# Tubular obstruction induced polycystin upregulation is pro-fibrotic and induced a severe cystic phenotype in adult mice with autosomal dominant polycystic kidney disease: the coexistence of polycystin loss and gain function in ADPKD

**DOI:** 10.1101/2021.11.03.467035

**Authors:** Ming Wu, Yanzhe Wang, Ying Jing, Dongping Chen, Yufeng Xing, Yanfang Bai, Di Huang, Yijing Zhou, Jinghua Hu, Shougang Zhuang, Chaoyang Ye

## Abstract

Mutations in *PKD1* (encoding polycystin-1) or *PKD2* (encoding polycystin-2) gene cause autosomal dominant polycystic kidney disease (ADPKD), however high levels of polycystins are detected in renal tissues of ADPKD patients. Animal studies showed that loss and gain of function of polycystins are both pathogenic and can induce cystic phenotype in the kidney, which are associated with enhanced renal fibrosis. Recent studies showed that increased expression of polycystins contributes to organ fibrosis. However, the role of polycystins in renal tubulointerstitial fibrosis remains unclear. In this study, we demonstrated that polycystin-1 or polycystin-2 was highly expressed in the kidney of two different fibrotic mouse models and positively correlated with expression of collagen-I. Pharmaceutical inhibition of polycystin-2 with triptolide or genetic knockout of polycystin-2 reduced the expression of epithelial-mesenchymal transition (EMT) markers and deposition of extracellular matrix proteins in fibrotic kidneys. Similarly, conditional knockout of *Pkd1* gene also attenuated renal fibrosis in mouse models. Thus, we further hypothesized that inhibition of polycystins delays cyst growth by mitigating renal fibrosis. Here, we showed that polycystin-1 or polycystin-2 was up-regulated in *Pkd2* or *Pkd1* mice respectively and tightly correlated with the growth of renal cysts and fibrosis development. Genetic deletion of both polycystin-1 and polycystin-2 retarded cyst growth in *Pkd1* or *Pkd2* mice. Finally, we deleted pkd1 gene in a fibrosis triggered adult ADPKD mouse model at different time point before or after the fibrotic injury. We showed that early and long-term inactivation of *Pkd1* delayed fibrosis triggered renal cyst growth in adult *Pkd1* mice as compared with mice with late and short-term inactivation of *Pkd1* gene. We conclude that tubular obstruction induced polycystin up-regulation is pro-fibrotic and accelerates cyst growth through enhancing renal interstitial fibrosis in ADPKD mice. Our study indicates that ADPKD is caused by both loss and gain function of polycystins. Reduction of the aberrant upregulation of polycystins in cystic kidneys is a therapeutic option for ADPKD patients.

**Research highlights:**

- Polycystin1 and polycystin-2 are up-regulated in fibrotic kidneys
- Inhibition or deletion of polycystins inhibits EMT and attenuates renal tubulointerstitial fibrosis
- Upregulation of polycystin1 or polycystin-2 is positively correlated with fibrosis progression and renal cyst growth in ADPKD mice
- Double knockout of Pkd1 and Pkd2 gene inhibits renal cyst growth in ADPKD mice
- Long-term deletion of Pkd1 gene delayed fibrosis triggered renal cyst growth in ADPKD mice

## Introduction

Chronic kidney disease (CKD) is a worldwide health concern affecting more than 10% of global population with a high mortality and morbidity (1). Renal fibrosis is the common pathway of various CKD progressing to end stage renal disease regardless of its original causes (1). Hallmarks of renal fibrosis include excessive deposition of extracellular matrix (ECM) proteins (such as collagen-I, and fibronectin), over-expression of epithelial-mesenchymal transition (EMT) markers (such as N-cadherin, α-SMA and vimentin), activation of pro-fibrotic signaling pathways (such as the TGFβ/Smad3 signaling pathway) (2). Recent studies showed epigenetic alterations during the progression of renal fibrosis (3).

Polycystins were originally identified in autosomal dominant polycystic kidney disease (ADPKD), where mutations in *PKD1* (encoding polycystin-1) or *PKD2* (encoding polycystin-2) gene cause this disease (4). Deletion of *Pkd1 or Pkd2* gene in mice can reproduce the cystic phenotype in the kidney, which indicates that loss of function of polycystins is pathogenic (5). Interestingly, several studies showed that exogeneous overexpression of polycystin-1 induced cyst formation in the mouse kidney, indicating that gain of function of polycystin-1 is also pathogenic (6–8). Abnormal renal fibrosis is observed in advanced ADPKD patients and mice with *Pkd* gene deletion, however transgenic mice with polycystin-1 overexpression also exhibit enhanced renal fibrosis (6, 9, 10). The direct effect of polycystins on renal fibrosis remain unknown.

In this study, we aimed to investigate the role of polycystins in renal tubulointerstitial fibrosis and their underlying mechanisms in models of renal fibrosis induced by unilateral ureteral obstruction (UUO) and aristolochic acid using genetic and pharmacological approaches.

## Results

### The expression of polycystin-2 is increased in two mouse models of renal tubulointerstitial fibrosis

As the first step towards understanding the role of polycystins in renal fibrosis, we examined polycystin-2 expression in two mouse models of renal tubulointerstitial fibrosis in wild type (WT) induced by UUO and aristolochic acid I. Compared with sham kidneys, the expression of polycystin-2 was steadily increased from day 3 to 14 in the kidney following UUO, which was correlated with the expression of collagen-I over time (Figure 1A-1C). In the model of aristolochic acid nephropathy (AAN), polycystin-2 and collagen 1 began to increase at 3 days and further elevated at day 21 following aristolochic acid administration (Figure 1D-1F).

**Fig. 1.**
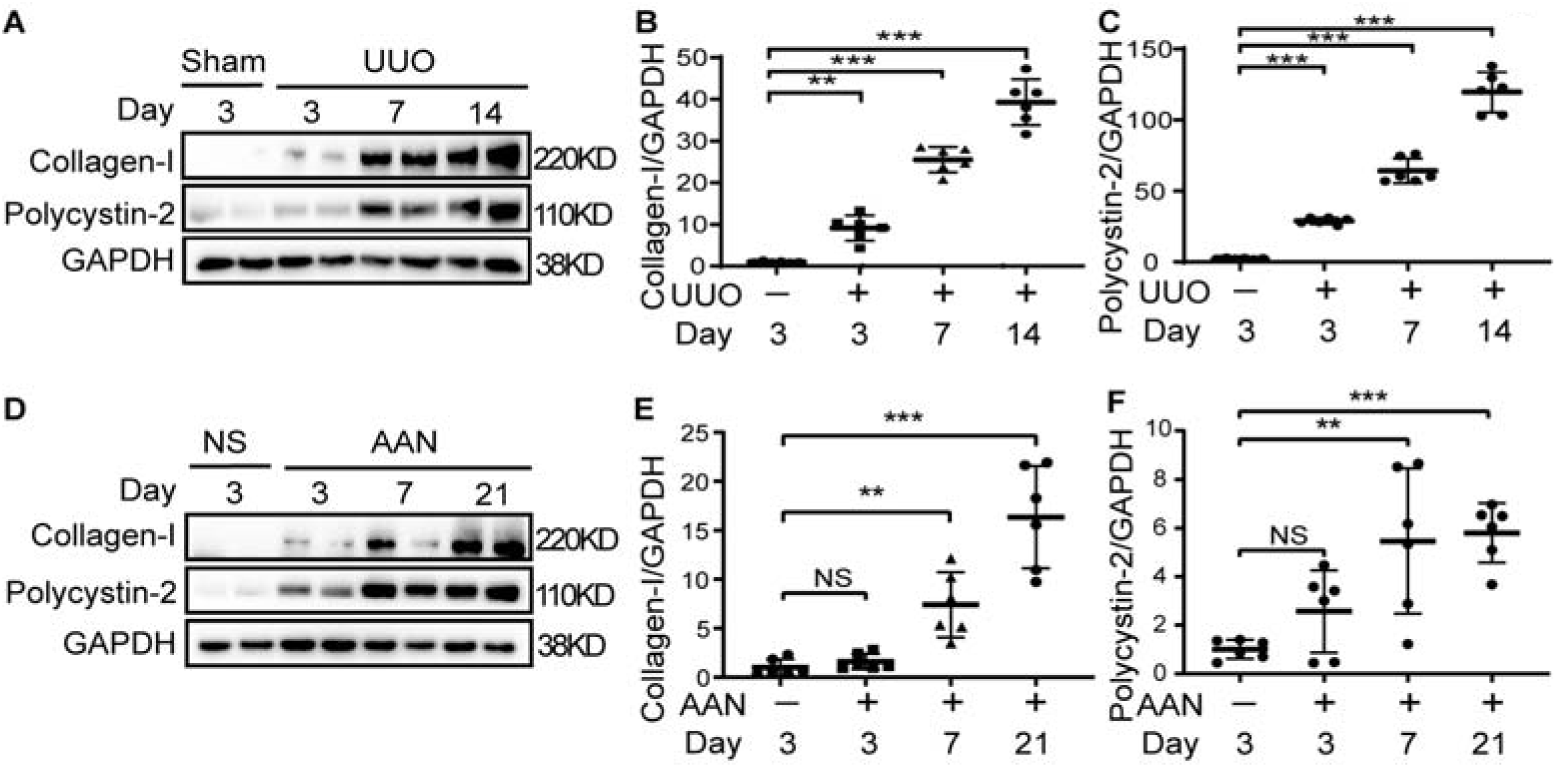
Polycystin-2 is up-regulated in fibrotic kidneys. (A-C) The time course of polycystin-2 expression in sham or unilateral ureteral obstruction (UUO) kidneys from wide type (WT) mice. The expression of collagen-I and polycystin-2 was analyzed by Western blotting and quantified. One representative of at least three independent experiments is shown. (D-F) The time course of polycystin-2 expression in aristolochic acid nephropathy (AAN) kidneys from WT mice. The expression of collagen-I and polycystin-2 in normal saline (NS) or AAN kidneys was analyzed by Western blotting and quantified. *N*[=5-6 in each experimental group and one representative of at least three independent experiments is shown. NS represents not significant. \*\**p*□<□0.01. \*\*\**p*□<□0.001.

### Inhibition of polycystin-2 attenuates renal tubulointerstitial fibrosis

Previous studies have shown that triptolide can bind to polycystin-2 and regulate its functions (11), we attempted to use triptolide as a molecular probe to examine the role of polycystin-2 in renal tubulointerstitial fibrosis in WT UUO mice. Masson staining showed that a massive deposition of interstitial collagens in UUO kidneys, and treatment with triptolide largely reduced extracellular matrix deposition (Figure 2A and 2B). Immunoblot analysis also indicated increased expression of ECM proteins (collagen-I) and EMT marker (α-SMA), and activated Smad3 (phosphorylated Smad3, pSmad3) in UUO kidneys and triptolide administration reduced expression of all those pro-fibrotic proteins (Figure 2C, and 2E-2G). Notably, triptolide also inhibited expression of polycystin-2 in kidneys following UUO operation (Figure 2D and 2H). These data illustrate that triptolide protected kidneys against renal fibrosis and suggest that polycystin-2 upregulation is required for the development of renal fibrosis.

**Fig. 2.**
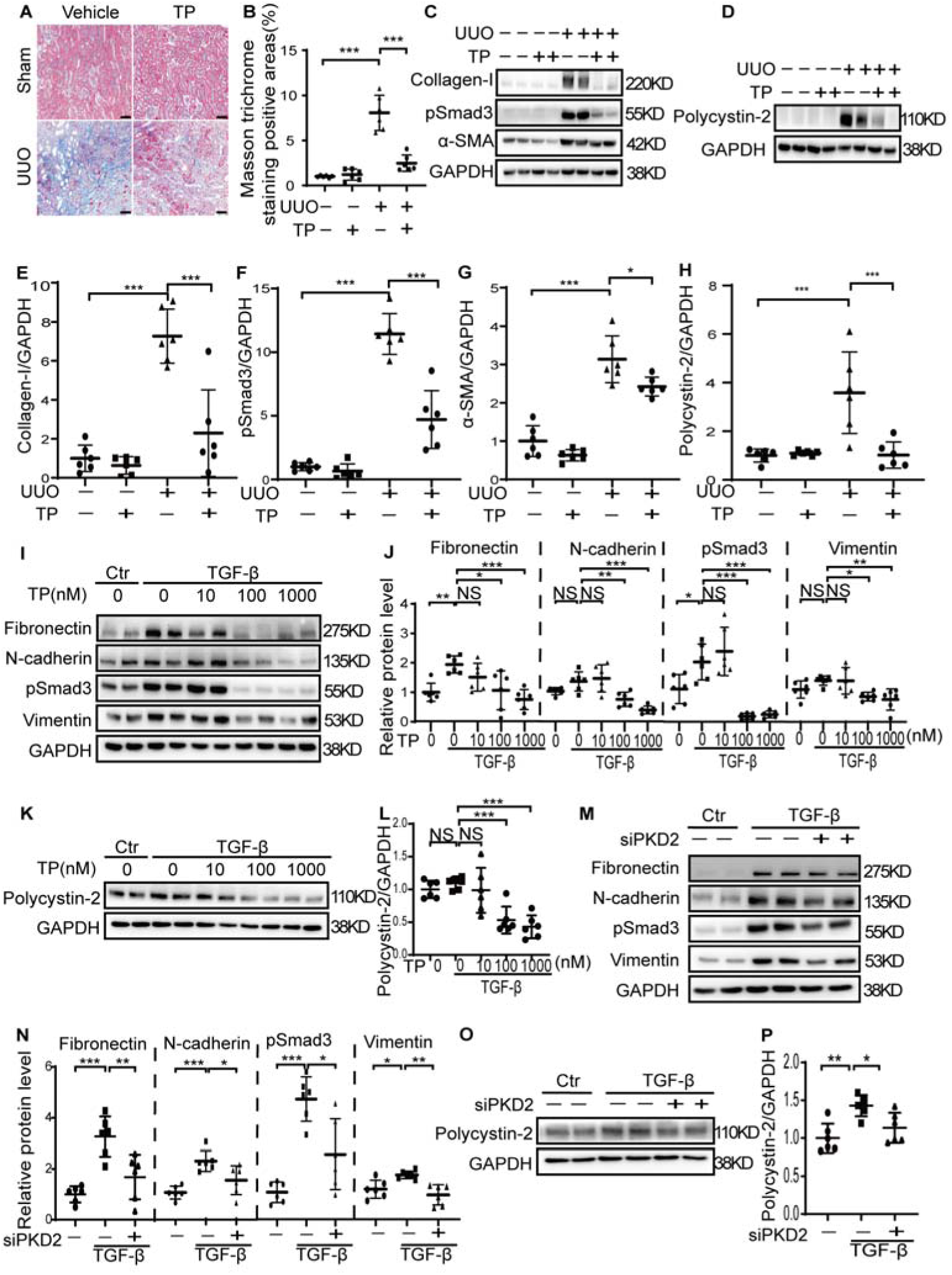
Pharmaceutical or genetic inhibition of polycystin-2 attenuates renal tubulointerstitial fibrosis. WT mice received sham or UUO operation followed by treatment with vehicle or triptolide (TP) for 10 days and then were sacrificed at day 10. (A-B) Renal fibrosis was assessed by Masson’s trichrome staining and quantified. Scale bar represents 100 μm. (C-H) The expression of collagen-I, phosphorylated Smad3 (pSmad3), α-SMA and polycystin-2 was analyzed by Western blotting and quantified. (I-L) HK2 human renal epithelial cells were stimulated with TGF-β and treated with different concentration of TP for 48 hours. Cell lysates were collected, and Western blot analysis was performed to determine expression of polycystin-2, fibronectin, N-cadherin, vimentin, and pSmad3, which was further quantified. (M-P) HK2 cells were transfected with nonsense control (NC) or PKD2 siRNA (siPKD2). On the second day, HK2 cells were stimulated with TGF-β for 48 hours. Cell lysates were collected, and Western blot analysis was performed to determine expression of polycystin-2, fibronectin, N-cadherin, vimentin, and pSmad3. *N*[=7-9 in each experimental group and one representative of at least three independent experiments is shown. NS represents not significant. \**p*□<□0.05. \*\**p*□<□0.01. \*\*\**p*□<□0.001.

To verify the role of polycystin-2 in renal fibrosis, we examined the effect of Triptolide and polycystin-2 siRNA on the expression of EMT associated proteins in cultured human renal epithelial (HK2) cells. As shown in Figure 2I-2J, exposure of HK2 cells to TGF-β1 increased expression of fibronectin, N-cadherin and vimentin, and pSmad3, and presence of triptolide from 100 nM to 1000 nM reduced their expression (Figure 2I-2J), which is corresponding to its inhibition on polycystin-2 expression (Figure 2K and 2L). Similarly, transfection of PKD2 siRNA also reduced expression of these pro-fibrotic proteins, which was correlated with down-regulation of polycystin-2 (Figure 2M and 2P).

Since it has been reported that polycystin-2 is expressed in tissue fibroblasts (12), we also examined the effect of triptolide on the expression of proteins as described above in cultured normal rat kidney fibroblasts (NRK-49F). In line with our observations in HK2 cells, triptolide dose-dependently inhibited expression of these pro-fibrotic markers and polycystin-2 in this cell line (supplemental Figure 1A-1F).

Collectively, these data suggest that injury-induced upregulation of polycystin-2 may contribute to EMT, fibroblast activation and renal fibrosis.

### Conditional knockout of polycystin-2 mitigates renal tubulointerstitial fibrosis in **mice**

To confirm the pro-fibrotic role of polycystin-2, we generated *Pkd2* conditional knockout mice. As shown in Figure 3A and 3B, heterozygous (*Pkd2^fl/+^;Cre/Esr1^+^*) or homozygous (*Pkd2^fl/fl^;Cre/Esr1^+^*) conditional knockout mice of *Pkd2* exhibited reduced positive areas of Masson staining in UUO kidneys. Although the similar degree of dilated tubules was present in control (*Pkd2^fl/+^;Cre/Esr1^-^*) or heterozygous *Pkd2* knockout UUO kidneys without appearance of cysts, numerous small cysts (luminal diameter exceeding 50 μm) in addition of dilated tubules were observed in the kidney of homozygous *Pkd2* knockout mice (Figure 3A). Immunoblot analysis showed partial reduction of *Pkd2* expression in heterozygous and its absence in homozygous conditional knockout of *Pkd2*, which is corresponding to a different degree of pro-fibrotic markers (collagen-I, pSmad3, and α-SMA) (Figure 3C-3G).

**Fig. 3.**
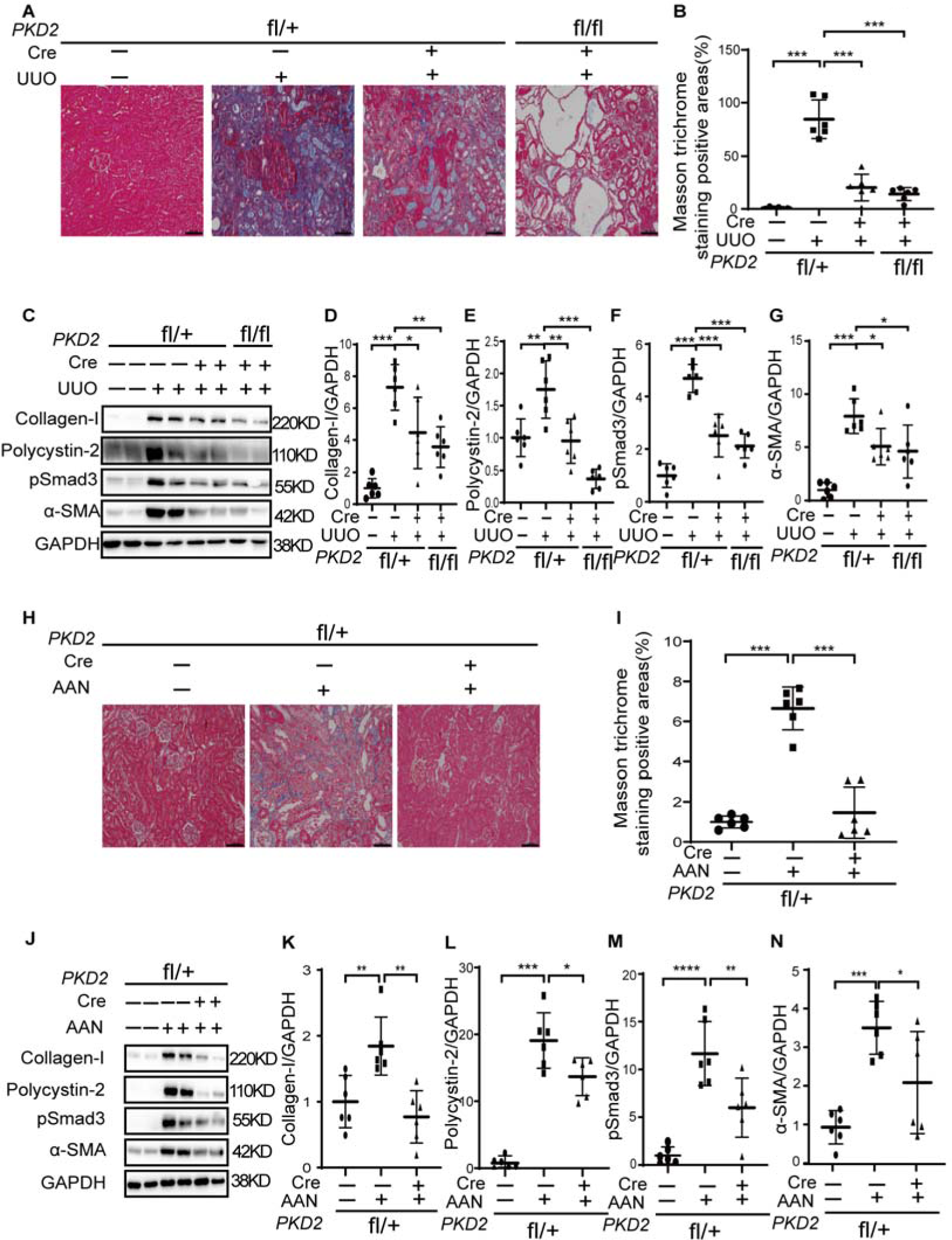
Conditional knockout of *Pkd2* attenuates renal tubulointerstitial fibrosis in mice. Control (*Pkd2^fl/+^;Cre/Esr1^-^*), heterozygous (*Pkd2^fl/+^;Cre/Esr1^+^*) or homozygous (*Pkd2^fl/fl^;Cre/Esr1^+^*) *Pkd2* conditional knockout mice received sham or UUO operation and were sacrificed at day 14. (A-B) Renal fibrosis was assessed by Masson’s trichrome staining. Scale bar represents 100 μm. (C-G) The expression of collagen-I, pSmad3, α-SMA and polycystin-2 was analyzed by Western blotting. To induce aristolochic acid nephropathy (AAN), control (*Pkd2^fl/+^;Cre/Esr1^-^*) or heterozygous (*Pkd2^fl/+^;Cre/Esr1^+^*) *Pkd2* conditional knockout mice were injected intraperitoneally with aristolochic acid I (AAI) or normal saline (NS) once a week for 2 weeks. *Pkd2* gene was inactivated at one week before AAI or NS injection. Mice were sacrificed at 3 weeks after the second AAI or NS injection. (H and J) Renal fibrosis was assessed by Masson’s trichrome staining and quantified. Scale bar represents 100 μm. (I, K, L, M) The expression of collagen-I, pSmad3, α-SMA and polycystin-2 was analyzed by Western blotting and further quantified. *N*□=6-8 in each experimental group and one representative of at least three independent experiments is shown. \**p*□<□0.05. \*\**p*□<□0.01. \*\*\**p*□<□0.001.

The anti-fibrotic effect of polycystin-2 deletion was observed in AAN mouse models as well. As shown in Figure 3H-3N, mild renal interstitial fibrosis was seen in control (*Pkd2^fl/+^;Cre/Esr1^-^*) AAN kidneys, as indicated by Masson staining and immunoblot analysis of collagen-I, pSmad3, and α-SMA (Figure 3H-3N). Heterozygous deletion of *Pkd2* gene in *Pkd2^fl/+^;Cre/Esr1^+^*mice showed reduced deposition of extracellular matrix and expression of pro-fibrotic markers in AAN kidneys, which was correlated with downregulation of polycystin-2 (Figure 3H-3N).

Together, all these data indicate that polycystin-2 promotes EMT and is pro-fibrotic in diseased kidneys.

### Polycystin-1 is increased in fibrotic kidneys and conditional deletion of *Pkd1* gene attenuates renal tubulointerstitial fibrosis in mouse kidneys

Polycystin-2 intertwined with polycystin-1 in primary cilia mediates all the biological responses, we thus further examined the expression of polycystin-1 in the mouse model with renal tubulointerstitial fibrosis. Compared with sham kidneys, the expression of polycystin-1 was increased from day 3 to day 14 in UUO kidneys (Figure 4A-4C).

**Fig. 4.**
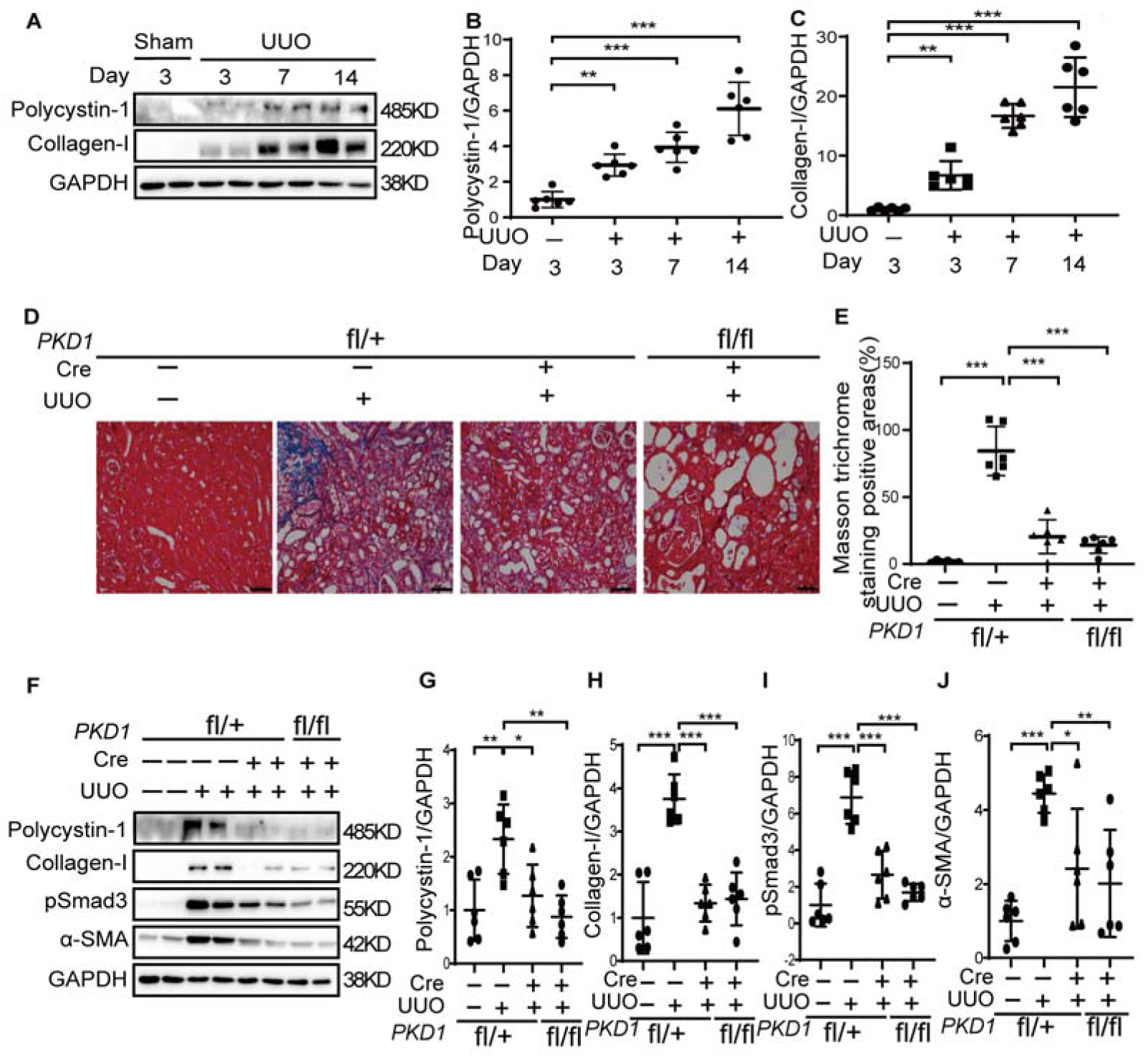
Conditional knockout of *Pkd1* attenuates renal tubulointerstitial fibrosis in injured kidneys induced by UUO operation. (A-C) The time course of polycystin-1 expression in sham or UUO kidneys from WT mice. The expression of collagen-I and polycystin-1 was analyzed by Western blotting and quantified. *N*□=5-6 in each experimental group and one representative of at least three independent experiments is shown. Control (*Pkd1^fl/+^;Cre/Esr1^-^*), heterozygous (*Pkd1^fl/+^;Cre/Esr1^+^*) or homozygous (*Pkd1^fl/fl^;Cre/Esr1^+^*) *Pkd1* knockout mice received sham or UUO operation and were sacrificed at day 14. (D-E) Renal fibrosis was assessed by Masson’s trichrome staining. Scale bar represents 100 μm. (F-J) The expression of collagen-I, pSmad3, α-SMA and polycystin-1 was analyzed by Western blotting and quantified. One representative of at least three independent experiments is shown. *N*□=6-8 in each experimental group and one representative of at least three independent experiments is shown. \**p*□<□0.05. \*\**p*□<□0.01. \*\*\**p*□<□0.001.

Using *Pkd1* conditional knockout mice, we found that heterozygous (*Pkd1^fl/+^;Cre/Esr1^+^*) or homozygous (*Pkd1^fl/fl^;Cre/Esr1^+^*) conditional knockout of *Pkd1* gene reduced the deposition of extracellular matrix in UUO kidneys as shown by Masson staining (Figure 4D-4E). Numerous small cysts were observed in homozygous *Pkd1* knockout mice (Figure 4D). Heterozygous or homozygous conditional knockout of *Pkd1* inhibited expression of polycystin-1 and pro-fibrotic markers (collagen-I, pSmad3, and α-SMA) in UUO kidneys as compared with control (*Pkd1^fl/+^;Cre/Esr1^-^*) UUO kidneys (Figure 4F-4J). Collectively, our data indicate that polycystin-1 promotes renal tubulointerstitial fibrosis as well.

### Genetic deletion of polycystin-2 delays disease progression in *Pkd1* mice

ADPKD is characterized by enhanced renal tubulointerstitial fibrosis, and thus we further hypothesized that deletion of polycystins may attenuate renal fibrosis and delay ADPKD progression. *Pkd1* or *Pkd2* gene inactivation was induced by peritoneally injection of tamoxifen from postnatal day 25 to 28 (P25-P28), and mice were sacrificed at P140 or P90 respectively (Figure 5A). Massive renal cysts were observed in *Pkd1* mice (*Pkd1^fl/fl^;Cre/Esr1^+^*) at P140 and in *Pkd2* mice (*Pkd2^fl/fl^;Cre/Esr1^+^*) at P90 (Figure 5B). *Pkd2* mice were further crossed with *Pkd1* mice. We found that heterozygous deletion of *Pkd2* (*Pkd2^fl/-^;Pkd1^fl/fl^;Cre/Esr1^+^*) gene inhibited cyst growth in *Pkd1* mice (Figure 5B). Two kidney weight/total body weight (2KW/TBW) ratio, blood urea nitrogen (BUN) and creatinine (CREA) levels were assessed (Figure 5C). *Pkd2* mice has more severe phenotype compared to *Pkd1* mice in this study, and we only observed significant increase in 2KW/TBW ratio but not in BUN or CREA levels in *Pkd2* mice as compared with that in control mice (*Pkd1^fl/fl^;Cre/Esr1^-^*) (Figure 5C). The 2KW/TBW ratio in *Pkd2^fl/-^;Pkd1^fl/fl^;Cre/Esr1^+^*mice was not significant lower than that in *Pkd1* mice but lower than that in *Pkd2* mice (Figure 5C).

**Fig. 5.**
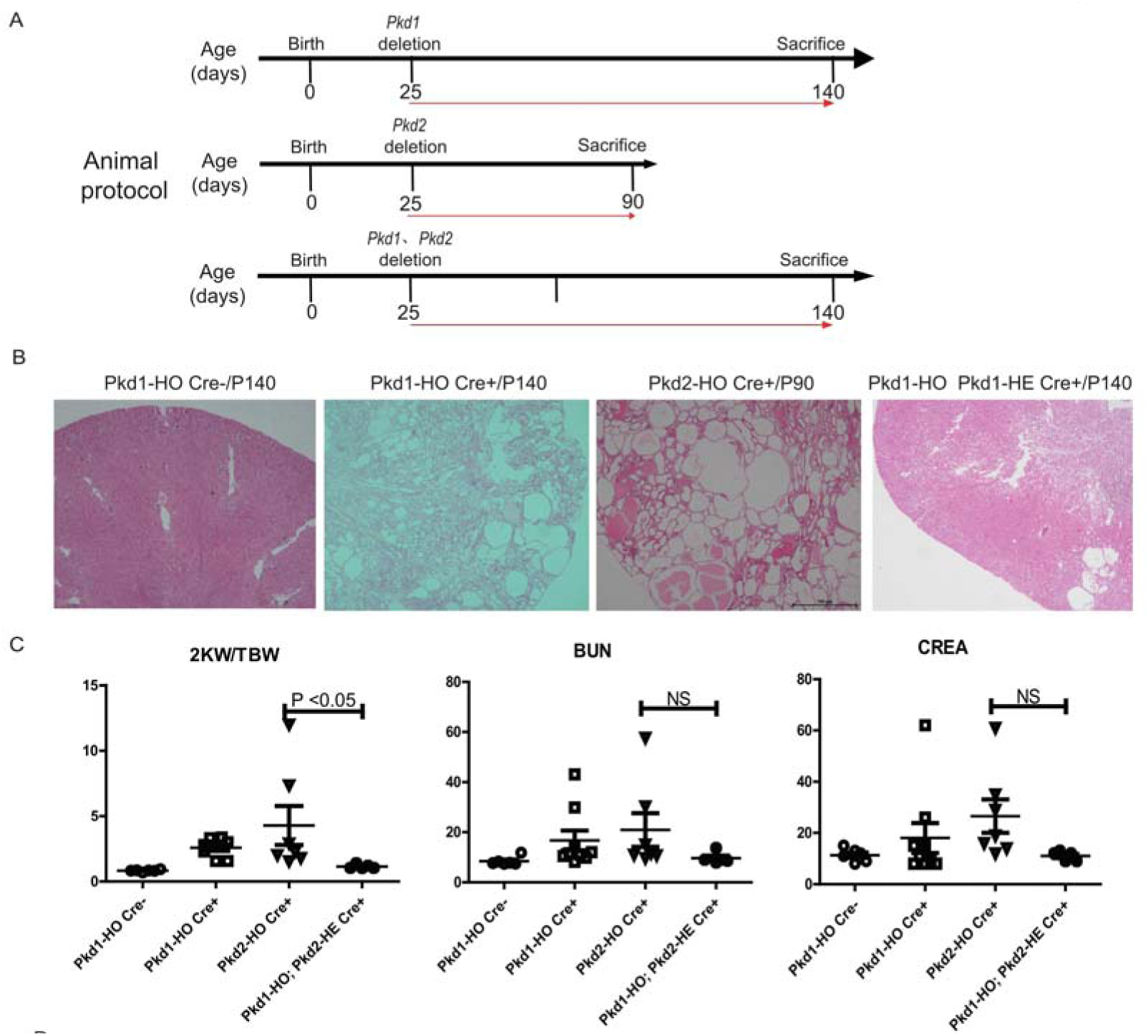
Deletion of polycystin retards disease progression in ADPKD mice. (A). *Pkd1* or *Pkd2* gene was inactivated at postnatal day 25 (P25) by intraperitoneal injection of tamoxifen in male mice, which were sacrificed at P140 or P90 respectively. (B-C). Renal function, kidney weight, and cyst volume were assessed. NS represents not significant. \**p*□<□0.05.

### Long-term inactivation of *Pkd1* delayed fibrosis triggered renal cyst growth in adult *Pkd1* mice

Renal injury and inactivation of both allele of ADPKD genes are third hits required for cystogenesis in adult ADPKD mice (13, 14). We inactivated *Pkd1* gene in adult (8 weeks of age) *Pkd1* mice at one week before aristolochic acid I (AAI, 5 mg/kg body weight) injection (13wk model) and sacrificed mice at 13 weeks after AAI injection (13 weeks after three hits) (Figure 6A). In another protocol (3+10wk model), *Pkd1* gene was inactivated in adult *Pkd1^fl/fl^;Cre/Esr1^+^*mice at three weeks after AAI injection, and mice were sacrificed at 10 weeks after *Pkd1* gene inactivation (10 weeks after three hits) (Figure 6A). HE staining showed that no renal cyst was observed in normal saline treated control *Pkd1^fl/fl^;Cre/Esr1^+^* mice (Figure 6B). Numerous small renal cysts were observed in mice with 13 weeks long-term *Pkd1* inactivation, and surprisingly more cysts were formed in *Pkd1* mice with short-term gene inactivation (3+10wk model) (Figure 6B). Compared with normal saline treated control *Pkd1^fl/fl^;Cre/Esr1^+^* mice, kidney weight but not BUN or CREA levels were increased in mice with 13 weeks long-term *Pkd1* inactivation, which were further increased in *Pkd1* mice with short-term gene inactivation (Figure 6C).

**Fig. 6.**
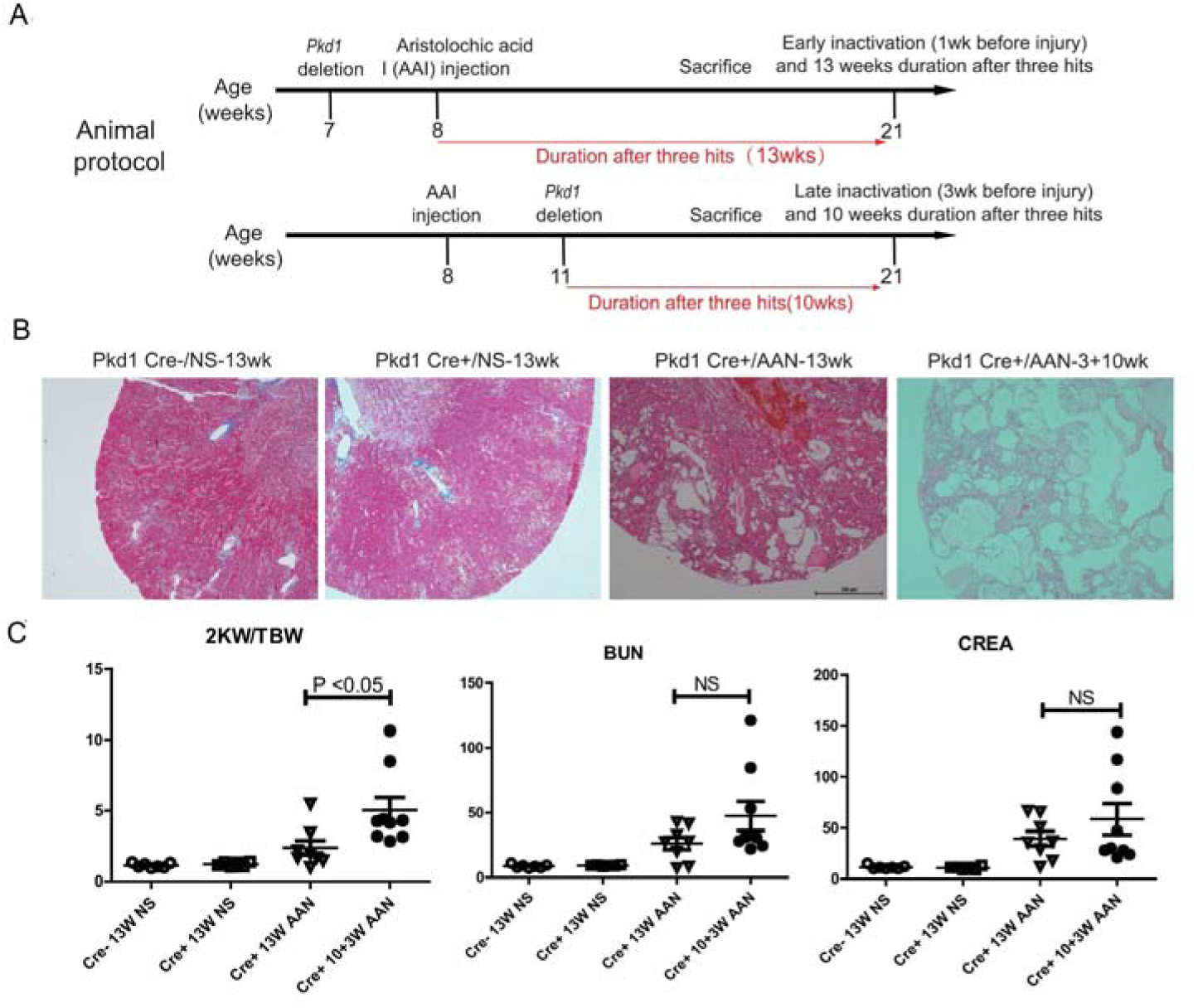
Long-term deletion of polycystin delays disease progression in fibrosis triggered adult ADPKD mice. (A). Renal injury was established by injection of AAI in male *Pkd1* mice at week 8, and Pkd1 gene was inactivated at one week before or three weeks after the kidney injury (three hits). Mice were sacrificed at 13 weeks after the kidney injury. (E-F). Renal function, kidney weight, and cyst volume were assessed. NS represents not significant. \**p*□<□0.05.

## Discussion

In the current study, we investigated the role of polycystins in renal tubulointerstitial fibrosis. We conclude that polycystins are pro-fibrotic in injured kidneys, which is supported by the following observations: (1) polycystin-1 and polycystin-2 were up-regulated in two different mouse models of renal fibrosis, which was positively correlated with the expression of ECM protein collagen-I; (2) inhibition of polycystin-2 by triptolide or siRNA reduced expression of pro-fibrotic markers in cultured renal epithelial cells; (3) pharmaceutical or genetic inhibition of polycystin-2 reduced partial EMT and renal fibrosis in UUO and AAN mouse models; (4) conditional knockout of *Pkd1* gene reduced pro-fibrotic marker expression and ECM accumulation in fibrotic kidneys.

Polcystin-1 and polycystin-2 form a transmembrane complex and are located in the primary cilia functioning as a calcium channel, which can sense external stimuli (12). Recent studies showed that primary cilia play an important role in organ fibrosis (15). Conditional knockout of ciliary proteins such as IFT88 or polycystin-1 attenuates fibrosis in injured lungs or hearts (12, 16). In line with these studies, inhibition or conditional knockout of the functional ciliary protein prolycystin-1 or polycystin-2 in injured kidneys also attenuated renal tubulointerstitial fibrosis, implying that primary cilia are required for renal tubulointerstitial fibrosis. Targeting ciliary genes or primary cilium related signaling pathways may be a new strategy to prevent or treat renal tubulointerstitial fibrosis.

Previous studies showed that renal fibrosis is increased in *Pkd1* or *Pkd2* deleted kidneys after renal ischemia reperfusion injuries (IRI) (17, 18). Nevertheless, the genetic animals and animal models in these two studies were different from ours. In these two studies, *Pkd1* or *Pkd2* gene was globally knockout, which led to a different baseline before the surgery, such as an increased cell proliferation in the kidney of knockout mice as compared with that in wide type mice (17, 18). We used conditional knockout mice in this study, and to keep the same baseline condition, we only inactivated *Pkd* gene at one week before kidney injury. Second, we used UUO and AAN models in this study, which are two classical models to study renal tubulointerstitial fibrosis. The pathogenesis of renal tubulointerstitial fibrosis in UUO, AAN and IRI models are different, which may lead to different renal outcomes in investigating the role of polycystins in renal fibrosis (19–21).

Partial EMT is an important mechanism underlying renal tubulointerstitial fibrosis (22). Under this pathological condition, renal tubular epithelial cells express profibrotic markers but do not migrate into interstitial areas (22). In this study, we showed that pharmacological inhibition or genetic deletion of polycystins reduced expression of EMT markers such as α-SMA, N-cadherin and vimentin in fibrotic kidneys and in TGF-β stimulated human renal epithelial cells, suggesting that polycystins are engaged in EMT. EMT also plays an important role in cancer progression (23). It has been reported that the cancer risk in ADPKD recipient is lower than that in non-ADPKD recipient after renal transplantation, however the underlying biological mechanism is currently unknown (24). Thus, it would be interesting to study whether polycystin-mediated EMT is involved in the pro-tumor action of ADPKD genes.

ADPKD is caused by heterozygous loss of polycystins (25). Paradoxically, polycystins are upregulated in kidney tissues of ADPKD patients, and thus studies on the gain of function of polycystins in kidney diseases is also important to understand the pathogenesis of ADPKD (6, 8). Based on the finding that polycystin overexpression is pro-fibrotic in kidney disease, we further hypothesized that reduction of the aberrant upregulation of polycystins may delay ADPKD progression. Here we showed that heterozygous deletion of polycystin-2 inhibited cyst growth in *Pkd1* mice. Interestingly, we further showed that long-term inactivation of polycystin-1 delayed fibrotic injury induced cyst growth in adult *Pkd1* mice as compared with *Pkd1* mice with short-term inactivation of polycystin-1.

A recent study showed that re-expression of *Pkd* genes reversed the progression of cystic kidney development in mice, suggesting that recovery of the lost *PKD* gene expression to normal levels by genetic therapy is a promising strategy for human ADPKD patients (26). Our study indicates that aberrant up-regulation of polycystins is pro-cystic and reduction of *Pkd* gene dosage slows cyst growth in ADPKD mice. Thus, properly control *PKD* gene dosage by either re-expression of the losted *PKD* gene to normal levels or reduction of upregulated polycystins could reverse or delay ADPKD progression in patients.

We conclude that tubular obstruction induced polycystin up-regulation is pro-fibrotic and accelerates cyst growth through enhancing renal interstitial fibrosis in ADPKD mice. Our study indicates that ADPKD is caused by both loss and gain function of polycystins. Reduction of the aberrant upregulation of polycystins in cystic kidneys is a therapeutic option for ADPKD patients.

## Materials and Methods

### Animals

Only male mice were used in animal studies. Wide type (WT) C57BL/6 mice (SPF grade, 20-25g) were purchased from Shanghai SLAC Laboratory Animal Co., Ltd. *Pkd1^fl/fl^* mice were obtained from Prof. Rudolf Wüthrich in University of Zurich, which were originally from the lab of Professor Gregory Germino in Johns Hopkins University School of Medicine (27). C57/BL6 *Pkd1^fl/fl^* mice were crossed with C57/BL6 tamoxifen-Cre (B6. Cg-Tg [Cre/Esr1]) 5Amc/J mice (stock 004682; Jackson Laboratories) as described previously (28). *Pkd2* conditional knockout mice with exon 3 deletion have cystic phenotype in the kidney (29). CRISPR/Cas9 technology was used to generate a mouse with loxP sites flanking exon 3 of the Pkd2 gene by Shanghai Model Organisms Center. *Pkd2^fl/fl^* were crossed with tamoxifen-Cre mice to generate *Pkd2^f^ ^l/f^ ^l^;Cre/Esr1^+^* mice. Animals were housed in the animal facility of Shanghai University of Traditional Chinese Medicine according to local regulations and guidelines. Animal experiments were approved by the ethic committee of Shanghai University of Traditional Chinese Medicine (PZSHUTCM18102604).

### Genotyping

Genomic DNA was prepared from mouse tail by using animal genomic DNA quick extraction kit (D0065M, Beyotime Biotech, Nantong, China) as described in the manufacturer’s instructions. PCR genotyping was performed using the following primers: Pkd2-F, GTACACCTGTGCAGCACTCA; Pkd2-R, GGAGCTCATCATGCCGTAGG; CRE-F, ATTGCTGTC ACTTGGTCGTGGC; CRE-R, GGAAAATGCTTCTGTCCGTTTGC; Pkd1-F, CCTGCCTTGCTCTACTTTCC; PKD1-R, AGGGCTTTTCTTGCTGGTCT; Details regarding PCR conditions are available upon request.

### Experimental Animal Protocol

UUO operation was performed through twice ligation of the left ureter with 4-0 nylon sutures. WT animals were sacrificed at day 3, 7 or 14 in a time-course experiment (n=5-6 for each time point). For interventional study, WT mice were randomly divided into four groups: 1) Sham/Vehicle (n=6), 2) Sham/triptolide (n=7), 3) UUO/Vehicle (n=7), and 4) UUO/triptolide (n=7) group. The stock of triptolide (Topscience, T2179, Shanghai, China) was dissolved in DMSO and further diluted in normal saline as working solution. Sham or UUO mice were treated with vehicle or 0.25 mg/kg triptolide daily by intraperitoneal (i.p.) injection for 10 days starting from day 0. Mice were sacrificed at day 10. Kidney samples were obtained for Western blotting or histological examinations. For conditional knockout mice (n= 7-9 mice per group), Cre recombinase activity was induced by intraperitoneally injection of tamoxifen (50 mg kg^-1^ day^-1^, Apexbio, B5965) for three consecutive days at one week before UUO operations. Conditional knockout mice at 8 weeks of age were subjected to UUO operation and were sacrifice at day14.

To induce aristolochic acid nephropathy in mice, one dose of aristolochic acid I (AAI, A9451, Sigma; 5 mg/kg body weight) or normal saline (NS) was peritoneally injected. WT animals were sacrificed at day 3, 7 or 21 in a time-course experiment (n=5-6 for each time point). Kidney tissues were collected. For conditional knockout mice (n= 6-8 mice per group), Cre recombinase activity was induced at 7 weeks of age by intraperitoneally injection of tamoxifen for three consecutive days at one week before the first AAI injection (5 mg/kg body weight), which was followed by a second AAI injection after one week (5 mg/kg body weight). Mice were sacrificed at 21 days after the second AAI injection.

To induce cystogenesis, *Pkd1* or *Pkd2* gene was inactivated at postnatal day 25 (P25) by intraperitoneal injection of tamoxifen for three consecutive days in male mice, which were sacrificed at P140 or P90 respectively. In another experiment, adult *Pkd1* mice were injected with AAI (5 mg/kg body weight) once at 8 weeks of age, *Pkd1* gene was inactivated by intraperitoneal injection of tamoxifen for three consecutive days at one week before or three weeks after the AAI injection. Mice were sacrificed at 13 weeks after the AAI injection. Mouse kidneys and blood were collected for further analysis.

### Cell Culture

Normal rat kidney 49F (NRK-49F) cells, a rat kidney interstitial fibroblast cell line, were purchased from national infrastructure of cell line resource, Chinese academy of medical sciences. To study the anti-fibrotic effect of triptolide, NRK-49F cells were seeded in 6-well plates to 40–50% confluence and were starved with DMEM/F12 medium containing 0.5% fetal bovine serum (FBS) overnight before the experiment. The next day, cells were refreshed with DMEM/F12 medium with 0.5% FBS and then were exposed to different concentrations of triptolide (10 nM, 100 nM, and 1000 nM) for 24h. Protein was extracted from cell lysates for further analysis.

HK2 renal proximal tubular epithelial cells were obtained from the Cell Bank of Shanghai Institute of Biological Sciences (Chinese Academy of Science). HK2 cells were seeded in 6-well plate to 40-50% confluence, which were starved overnight with DMEM/F12 medium containing 0.5% fetal bovine serum. On the next day, fresh medium containing 0.5% fetal bovine serum was changed, and then cells were exposed to 2.5 ng/ml TGF-β (Peprotech, Rocky Hill, NJ, USA) for 48h in the presence of various concentration of triptolide.

Nonsense control (NC) or human PKD2 siRNA were transfected by Lipofectamine 2000 (11668-027; Invitrogen) in HK2 cells using DMEM/F12 medium containing 10% fetal bovine serum according to the manufacturer’s instruction. Protein was extracted from cell lysates at 48h after transfection to assess the efficacy of PKD2 siRNA (supplemental Figure 2). In another experiment, fresh medium containing 0.5% FBS was changed on the second day after transfection, and then cells were exposed to 2.5 ng/ml TGF-β (Peprotech, Rocky Hill, NJ, USA) for 48h with or without triptolide. The NC siRNA sequences were as follows: forward, 5′-UUCUCCGAACGUGUCACGUTT-3′; and reverse, 5′-ACGUGACACGUUCGGAGAATT-3′. The human PKD2 siRNA sequences were as follows: forward, 5′-GGACAUAGCUCCAGAAGGATT-3′; and reverse, 5′-UCCUUCUGGAGCUAUGUCCTT-3′.

### Hematoxylin-eosin and Masson’s Trichrome Staining

Mouse kidneys were fixed and embedded in paraffin. Four-μm-thick sections of paraffin-embedded kidney tissue was stained with hematoxylin, and then with ponceau red liquid dye acid complex, which was followed by incubation with phosphomolybdic acid solution. Finally, the tissue was stained with aniline blue liquid and acetic acid. Images were obtained with the use of a microscope (Nikon 80i, Tokyo, Japan).

### Western Blotting

Cellular or kidney protein was extracted by using lysis buffer purchased from Beyotime Biotech (P0013, Nantong, China). The protein concentration was measured by using BCA Protein Assay Kit (P0012S, Beyotime Biotech, Nantong, China). Protein samples were dissolved in 5x SDS-PAGE loading buffer (P0015L, Beyotime Biotech, Nantong, China), which were further subjected to SDS-PAGE gel electrophoresis. After electrophoresis, proteins were electro-transferred to a polyvinylidene difluoride membrane (Merck Millipore, Darmstadt, Germany), which was incubated in the blocking buffer (5% non-fat milk, 20mM Tris-HCl, 150mMNaCl, PH=8.0, 0.01%Tween 20) for 1 hour at room temperature and was followed by incubation with anti-polycystin-1 (1:500, sc-130554, SantaCruz), anti-polycystin-2 (1:1000, 19126-1-AP, Proteintech), anti-polycystin-2 (1:1000, A3625, ABclonal), anti-KIF3A (1:500, SC-376680, SantaCruz), anti-fibronectin (1:1000, ab23750, Abcam), anti-pSmad3 (1:1000, ET1609-41, HUABIO), anti-Collagen I (1:1000, ab260043, Abcam), anti-α-SMA (1:1000, ET1607-53, HUABIO), anti-N-cadherin (1:1000, sc-59887, Santa Cruz), anti-vimentin (1:1000, R1308-6, HUABIO), anti-GAPDH (1:5000, 60004-1-lg, Proteintech), or anti-α-tubulin (1:1000, AF0001, Byotime) antibodies overnight at 4□. Binding of the primary antibody was detected by an enhanced chemiluminescence method (SuperSignal™ West Femto, 34094, Thermo Fisher Scientific) using horseradish peroxidase-conjugated secondary antibodies (goat anti-rabbit IgG, 1:1000, A0208, Beyotime or goat anti-mouse IgG, 1:1000, A0216, Beyotime).

### Statistical Analysis

Statistical analyses were performed by one-way ANOVA followed by a Dunnett’s multiple comparisons test using GraphPad Prism version 5.0 (GraphPad, San Diego, CA). All data are expressed as means ± SD, and P < 0.05 was considered as statistically significant.

## Author Contributions

CY funded the project. MW, SZ, JH, YJ, CY, DC conceived and coordinated the study. MW, SZ and JH wrote the paper. YW conducted the *in vitro* experiments. MW, YW, YJ, JL, DC, BT, MY and DH performed the animal experiments. YW performed and analyzed the Western blotting. All authors reviewed the results and approved the final version of the manuscript.

---

**Supplemental Fig. 1.**
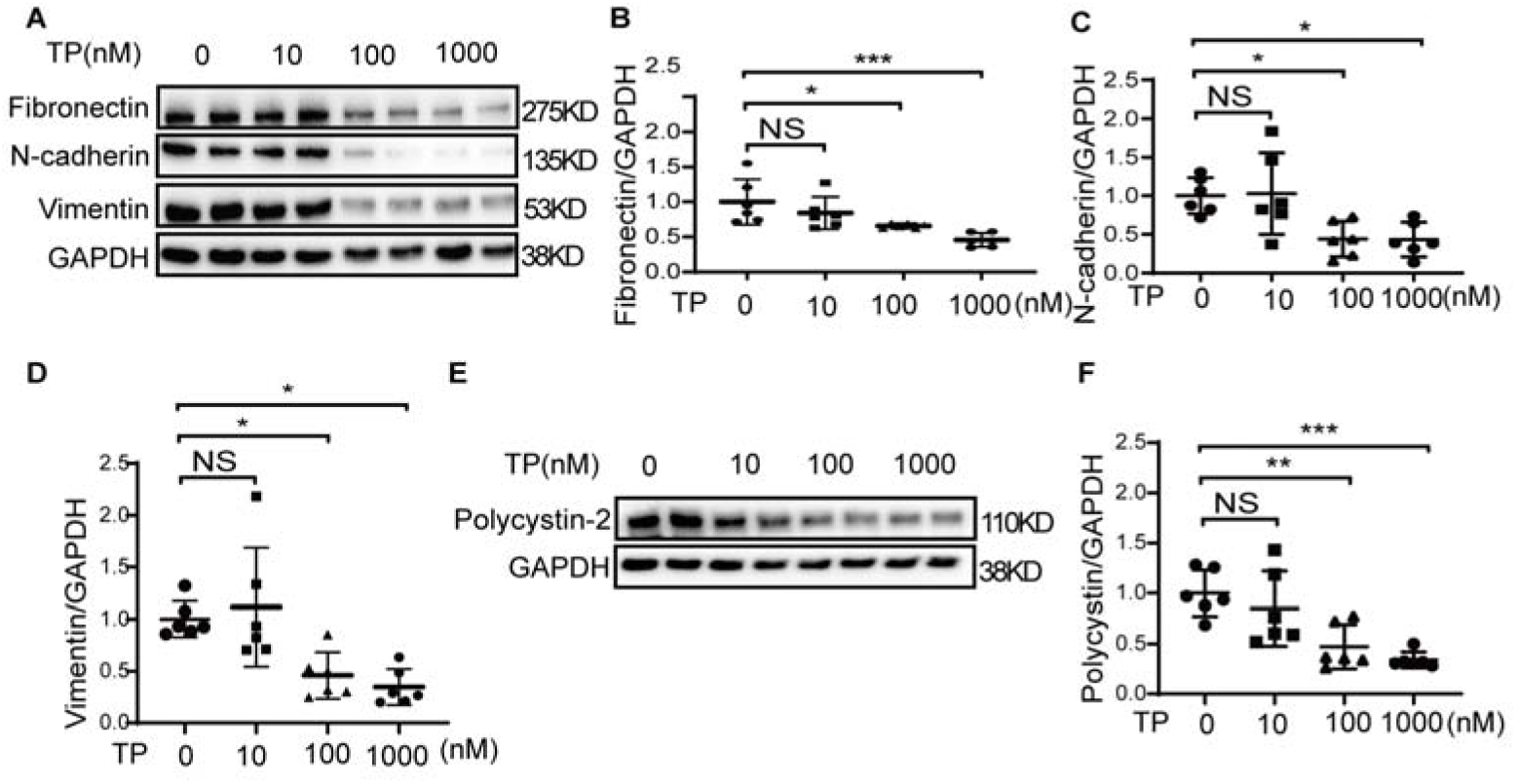
Triptolide reduces the expression of polycystin-2 and pro-fibrotic markers in renal fibroblasts. Normal rat renal fibroblasts (NRF-49F) were treated with different concentrations of triptolide (TP) for 24 hours, and cell lysates were collected. (A-F) Western blot analysis was performed to determine the expression of polycystin-2, fibronectin, N-cadherin, and vimentin, which was further quantified. One representative of at least three independent experiments is shown. NS represents not significant. \**p*□<□0.05. \*\**p*□<□0.01. \*\*\**p*□<□0.001.

**Supplemental Fig. 2.**
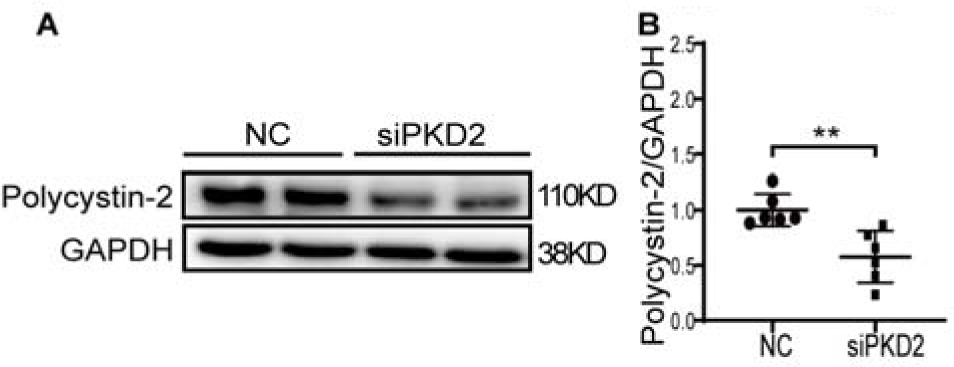
Transfection efficiency of PKD2 siRNA in HK2 cells. HK2 cells were transfected with NC or PKD2 siRNA. On the second day, HK2 cells were stimulated with TGF-β for 48 hours. Cell lysates were collected, and Western blot analysis and its quantification were performed to measure the expression of polycystin-2. One representative of at least three independent experiments is shown. \*\**p*□<□0.01.

## References

1. Gewin LS. Renal fibrosis: Primacy of the proximal tubule. Matrix Biol. 2018;68–69:248-62.

2. Schnaper HW. The Tubulointerstitial Pathophysiology of Progressive Kidney Disease. Adv Chronic Kidney Dis. 2017;24(2):107–16.

3. Li T, Yu C, and Zhuang S. Histone Methyltransferase EZH2: A Potential Therapeutic Target for Kidney Diseases. Front Physiol. 2021;12:640700.

4. Grantham JJ, Mulamalla S, and Swenson-Fields KI. Why kidneys fail in autosomal dominant polycystic kidney disease. Nat Rev Nephrol. 2011;7(10):556–66.

5. Ghata J, and Cowley BD, Jr. Polycystic Kidney Disease. Compr Physiol. 2017;7(3):945–75.

6. Kurbegovic A, Cote O, Couillard M, Ward CJ, Harris PC, and Trudel M. Pkd1 transgenic mice: adult model of polycystic kidney disease with extrarenal and renal phenotypes. Hum Mol Genet. 2010;19(7):1174–89.

7. Pritchard L, Sloane-Stanley JA, Sharpe JA, Aspinwall R, Lu W, Buckle V, et al. A human PKD1 transgene generates functional polycystin-1 in mice and is associated with a cystic phenotype. Hum Mol Genet. 2000;9(18):2617–27.

8. Thivierge C, Kurbegovic A, Couillard M, Guillaume R, Cote O, and Trudel M. Overexpression of PKD1 causes polycystic kidney disease. Mol Cell Biol. 2006;26(4):1538–48.

9. Mun H, and Park JH. Inflammation and Fibrosis in ADPKD. Adv Exp Med Biol. 2016;933:35–44.

10. Song CJ, Zimmerman KA, Henke SJ, and Yoder BK. Inflammation and Fibrosis in Polycystic Kidney Disease. Results Probl Cell Differ. 2017;60:323–44.

11. Leuenroth SJ, Okuhara D, Shotwell JD, Markowitz GS, Yu Z, Somlo S, et al. Triptolide is a traditional Chinese medicine-derived inhibitor of polycystic kidney disease. Proc Natl Acad Sci U S A. 2007;104(11):4389–94.

12. Villalobos E, Criollo A, Schiattarella GG, Altamirano F, French KM, May HI, et al. Fibroblast Primary Cilia Are Required for Cardiac Fibrosis. Circulation. 2019;139(20):2342–57.

13. Happe H, Leonhard WN, van der Wal A, van de Water B, Lantinga-van Leeuwen IS, Breuning MH, et al. Toxic tubular injury in kidneys from Pkd1-deletion mice accelerates cystogenesis accompanied by dysregulated planar cell polarity and canonical Wnt signaling pathways. Hum Mol Genet. 2009;18(14):2532–42.

14. Takakura A, Contrino L, Zhou X, Bonventre JV, Sun Y, Humphreys BD, et al. Renal injury is a third hit promoting rapid development of adult polycystic kidney disease. Hum Mol Genet. 2009;18(14):2523–31.

15. Bhattacharyya D, Teves ME, and Varga J. The dynamic organelle primary cilia: emerging roles in organ fibrosis. Curr Opin Rheumatol. 2021;33(6):495–504.

16. Kim E, Mathai SK, Stancil IT, Ma X, Hernandez-Gutierrez A, Becerra JN, et al. Aberrant Multiciliogenesis in Idiopathic Pulmonary Fibrosis. Am J Respir Cell Mol Biol. 2022;67(2):188–200.

17. Prasad S, McDaid JP, Tam FW, Haylor JL, and Ong AC. Pkd2 dosage influences cellular repair responses following ischemia-reperfusion injury. Am J Pathol. 2009;175(4):1493–503.

18. Bastos AP, Piontek K, Silva AM, Martini D, Menezes LF, Fonseca JM, et al. Pkd1 haploinsufficiency increases renal damage and induces microcyst formation following ischemia/reperfusion. J Am Soc Nephrol. 2009;20(11):2389–402.

19. Fu Y, Tang C, Cai J, Chen G, Zhang D, and Dong Z. Rodent models of AKI-CKD transition. Am J Physiol Renal Physiol. 2018;315(4):F1098–F106.

20. Chevalier RL, Forbes MS, and Thornhill BA. Ureteral obstruction as a model of renal interstitial fibrosis and obstructive nephropathy. Kidney Int. 2009;75(11):1145–52.

21. Zhou L, Fu P, Huang XR, Liu F, Chung AC, Lai KN, et al. Mechanism of chronic aristolochic acid nephropathy: role of Smad3. Am J Physiol Renal Physiol. 2010;298(4):F1006–17.

22. Sheng L, and Zhuang S. New Insights Into the Role and Mechanism of Partial Epithelial-Mesenchymal Transition in Kidney Fibrosis. Front Physiol. 2020;11:569322.

23. Saitoh M. Epithelial-Mesenchymal Transition by Synergy between Transforming Growth Factor-beta and Growth Factors in Cancer Progression. Diagnostics (Basel). 2022;12(9).

24. Wetmore JB, Calvet JP, Yu AS, Lynch CF, Wang CJ, Kasiske BL, et al. Polycystic kidney disease and cancer after renal transplantation. J Am Soc Nephrol. 2014;25(10):2335–41.

25. Capuano I, Buonanno P, Riccio E, Amicone M, and Pisani A. Therapeutic advances in ADPKD: the future awaits. J Nephrol. 2022;35(2):397–415.

26. Dong K, Zhang C, Tian X, Coman D, Hyder F, Ma M, et al. Renal plasticity revealed through reversal of polycystic kidney disease in mice. Nat Genet. 2021;53(12):1649–63.

27. Su Z, Wang X, Gao X, Liu Y, Pan C, Hu H, et al. Excessive activation of the alternative complement pathway in autosomal dominant polycystic kidney disease. J Intern Med. 2014;276(5):470–85.

28. Sun Y, Liu Z, Cao X, Lu Y, Mi Z, He C, et al. Activation of P-TEFb by cAMP-PKA signaling in autosomal dominant polycystic kidney disease. Sci Adv. 2019;5(6):eaaw3593.

29. Kim I, Ding T, Fu Y, Li C, Cui L, Li A, et al. Conditional mutation of Pkd2 causes cystogenesis and upregulates beta-catenin. J Am Soc Nephrol. 2009;20(12):2556–69.

